# Variation in filamentous growth and response to quorum-sensing compounds in environmental isolates of *Saccharomyces cerevisiae*

**DOI:** 10.1101/548099

**Authors:** B. Adam Lenhart, Brianna Meeks, Helen A. Murphy

## Abstract

In fungi, filamentous growth is a major developmental transition that occurs in response to environmental cues. In diploid *Saccharomyces cerevisiae*, it is known as pseudohyphal growth and presumed to be a foraging mechanism. Rather than normal unicellular growth, multicellular filaments composed of elongated, attached cells spread over and into surfaces. This morphogenetic switch can be induced through quorum sensing with the aromatic alcohols phenylethanol and tryptophol. Most research investigating pseudohyphal growth has been conducted in a single lab background, Σ1278b. To investigate the natural variation in this phenotype and its induction, we assayed the diverse 100-genomes collection of environmental *S. cerevisiae* isolates. Using computational image analysis, we quantified the production of pseudohyphae and observed a large amount of variation. Unlike ecological niche, population membership was associated with pseudohyphal growth, with the West African population having the most. Surprisingly, most strains showed little or no response to exogenous phenylethanol or tryptophol. We also investigated the amount of natural genetic variation in pseudohyphal growth using a mapping population derived from a single, highly-heterozygous clinical isolate that contained as much phenotypic variation as the environmental panel. A bulk-segregant analysis uncovered five major peaks with candidate loci that have been implicated in the Σ1278b background. Our results indicate that the filamentous growth response is a generalized, highly variable phenotype in natural populations, while response to quorum sensing molecules is surprisingly rare. These findings highlight the importance of coupling studies in tractable lab strains with natural isolates in order to understand the relevance and distribution of well-studied traits.

## Introduction

The budding yeast, *Saccharomyces cerevisiae*, can respond to environmental cues with numerous morphological switches and developmental phenotypes that likely increase fitness in naturally occurring conditions (Zaman et al. 2008). One such phenotype, filamentous growth, is thought to be a foraging strategy in response to nutrient stress. It is characterized by elongated cell morphology, unipolar budding, incomplete separation of mother-daughter cells, and substrate invasion (Gimeno et al. 1992). In diploid cells it is primarily induced by nitrogen limitation and known as pseudohyphal growth (Figure 1), while a similar though distinct response is triggered by carbon source limitation in haploid cells and is known as haploid invasive growth (Cullen and Sprague 2000, Cullen and Sprague 2012).

**Figure 1:**
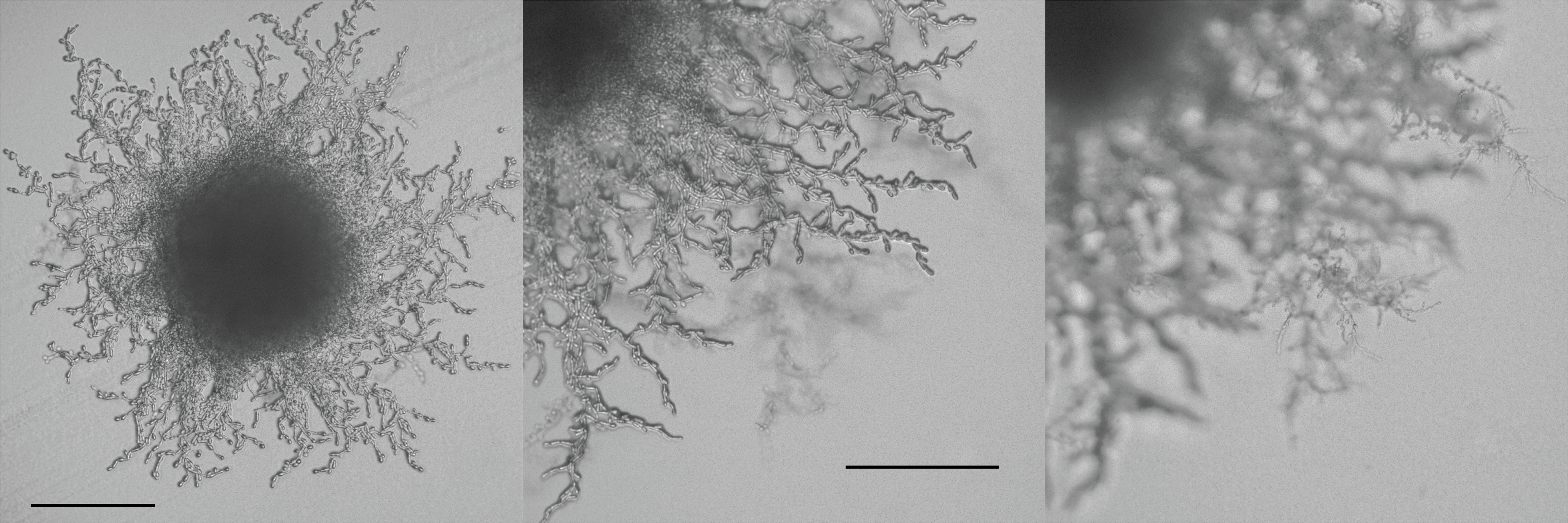
Pseudohyphal growth. Images depict: a small colony with pseudohyphae surrounding it (A), a close-up of a colony perimeter (B), and an image of the same perimeter in a different focal plane showing the pseudohyphae growing into the agar (C). Scale bar represents 200 um. To obtain images, strain YJM1439 was streaked on a SLAD plate and grown for 6 days.

In a lab strain, Σ1278b, haploid and diploid filamentous growth were shown to occur in response to the aromatic alcohols phenylethanol and tryptophol (Chen and Fink 2006). Production of these compounds is dependent on cell density and regulated through positive feedback, suggesting they may function as auto-inducing, quorum-sensing (QS) molecules. The human commensal and opportunistic pathogen, *Candida albicans*, can also undergo a morphological switch to a form of filamentous growth in response to QS molecules, which may be related to its ability to be pathogenic (Hornby et al. 2001, Leberer et al. 2001, Rocha et al. 2001, Chen et al. 2004, Biswas et al. 2007, Mallick and Bennett 2013). Parts of the signaling pathway are evolutionarily conserved (Lo et al. 1997, Cain et al. 2012); thus, filamentous growth may represent a general, social, yeast survival strategy in the natural environment (Wuster and Babu 2010).

In *S. cerevisiae*, filamentous growth is regulated by multiple evolutionarily conserved, pleiotropic signaling networks, including the glucose-sensing RAS/cAMP-PKA and SNF pathways, the nutrient-sensing TOR pathway, and the filamentous growth MAPK pathway (reviewed in (Granek et al. 2011, Cullen et al. 2012)). These signaling pathways converge to regulate the transcription of *FLO11*, which encodes a cell wall flocculin required for multiple *S. cerevisiae* developmental phenotypes, including filamentous growth (Lambrechts et al. 1996, Lo and Dranginis 1998, Pan and Heitman 1999, Rupp et al. 1999, Braus et al. 2003, Chen and Thorner 2010). In lab backgrounds, deletion (Jin et al. 2008, Ryan et al. 2012) and overexpression collections (Shively et al. 2013), as well as QTL mapping of genetic crosses (Song et al. 2014, Matsui et al. 2015), have identified hundreds of genes contributing to the phenotype.

The extent of phenotypic and genetic variation in filamentous growth in natural populations is still under-explored, as studies of this phenotype are dominated by the lab strain Σ1278b. Previous work has shown that in comparison to natural isolates, lab strains are often genetically and phenotypically atypical (Warringer et al. 2011). Thus, incorporating environmental isolates into genetic research broadens the scope of our understanding, both in how genetic variation modulates traits and how phenotypic variation manifests in natural populations (Gasch et al. 2016). The present study makes use of the 100-genomes collection, a panel of yeasts from subpopulations around the world and from a broad diversity of ecological niches, including fermentation reactions, clinical patients, and soil, plant and insect samples (Strope, et al., 2015), to explore natural variation in filamentous growth and response to QS compounds.

*Saccharomyces* yeasts are presumed diploid in nature (Replansky et al. 2008); therefore, the focus of the present study is the diploid filamentous growth response, pseudohyphal growth (psh). Most studies of psh use agar invasion as a quantitative metric for the phenotype, while the appearance of “pseudohyphae” around a colony (Figure 1) is assessed qualitatively. Using these metrics, (Magwene et al. 2011) found variation in a sample of environmental isolates, and Hope and Dunham (2014) found variation in the SGRP collection (Liti et al. 2009). Less is known about variation in psh response to QS molecules, and to our knowledge, no systematic surveys have been done.

Using a quantitative measure of the amount of pseudohyphae to estimate psh, which we call the “filamentous index”, we found a large amount of variation in the 100-genomes collection. When the strains were classified by the niche from which they were isolated, fermentation isolates exhibited a slightly elevated filamentous index compared to other ecological niches; however, this result appeared driven by a single isolate. When strains were classified by their subpopulation membership (Figure 2), which took phylogenetic history into consideration, isolates from the West African subpopulation had an elevated filamentous index compared to other subpopulations. Surprisingly, we find that in most isolates, addition of either phenylethanol or tryptophol to the medium had a negligible effect on psh, with a few strain-specific exceptions.

**Figure 2:**
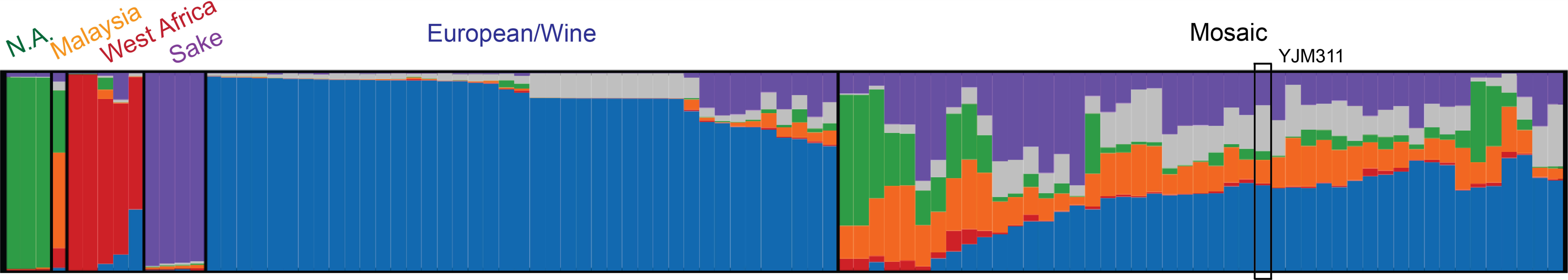
Population structure of the 100-genomes panel supplemented with strain YJM311, as inferred by the program *structure* (Pritchard et al. 2000) and following the analysis of Strope et al. (2015). Each vertical line represents an individual strain with its fractional ancestry of K=6 subpopulations represented by colors: green (North American), orange (Malaysian), red (West African), purple (Sake), blue (European/wine), and gray (human associated). Strains were assigned membership based on a threshold of >60% ancestry in a subpopulation, except for mosaic strains which had less than 60% ancestry in any other subpopulation.

As most genomic studies focus on Σ1278b, the present study also examined the amount of naturally occurring, segregating genetic variation for psh using a mapping population of segregants from YJM311 as a proxy. This strain is a highly-heterozygous clinical isolate (Granek et al. 2013) belonging to the “mosaic” subpopulation that contains genetic variation associated with each of the other major *S. cerevisiae* subpopulations (Figure 2). As such, it represents an ideal representative genetic background to investigate. We find that this single background contains enough genetic variation to recapitulate the range of phenotypes found in the 100-genomes panel. Using a bulk-segregant analysis, we find 5 genomic regions with major peaks significantly associated with the traits. Numerous genes that have been shown to influence the trait in Σ1278b are located within the peaks, and could therefore plausibly harbor the causative alleles.

Overall, our results indicate that there is an extensive amount of phenotypic and genetic variation in a well-studied developmental phenotype in environmental isolates, and that the response to aromatic alcohols may be a more limited, strain-specific effect. The relevance of this phenotype in the natural environment remains unknown, as no single ecological niche appeared to be strongly associated with the trait, while subpopulation membership did seem to be associated with psh ability. Our results highlight the importance of complementing studies in lab strains with numerous genetic backgrounds isolated from the environment.

## Methods and Materials

### Strains

96 strains from the 100-genomes collection (Strope et al. 2015) were phenotyped for psh (Table S1); these diploid strains are derived from single spores from original environmental isolates. Three of the strains, wells H8, H9, and H10 are not *S. cerevisiae*, and were not included the downstream analyses. YJM311, a homothallic, clinical isolate (Granek et al. 2013), was used to conduct a bulk segregant analysis (BSA). For a different study in our lab, the original diploid isolate was transformed to express a *PGK1*-*mCherry-KanMX* fusion (HMY7) and used to generate an F5 mapping population. This mapping population was used in the present study.

### Media

Yeast were grown in liquid YPD (1% yeast extract, 2% peptone, and 2% dextrose). Psh was induced on 4X synthetic low-ammonium dextrose (SLAD; 0.68% yeast nitrogen base w/o amino acids or ammonium sulfate, 2% dextrose, 50 µmol ammonium sulfate, and 2% agar) (Chen et al. 2006) and when appropriate, supplemented with phenylethanol (PheOH; Sigma-Aldrich, 77861) or Tryptophol (TrpOH; Sigma-Aldrich, T90301) dissolved in DMSO, added to a final concentration of 100 µmol. OmniTrays were poured two days before use in an assay. Sporulation of the mapping population was induced on sporulation medium (2% potassium acetate, 2% agar).

### Generation of an F5 Mapping Population

HMY7 was cycled through 4 rounds of sporulation, digestion, mating and growth (Supplementary Text). At the end of the last cycle, spores were plated to a density of ∼100 colonies per plate and 360 segregants were isolated and phenotyped. Each colony was presumed diploid due to self-mating.

### Sequencing and Bulk Segregant Analysis

Segregants with the highest and lowest filamentous indices, as well as lowest variance among replicate measurements, were chosen for further analysis. After re-assaying to verify psh, 22 segregants were identified for each pool. Segregants were grown to saturation in YPD in a 96-well plate, then combined for total genomic DNA extraction with the MasterPure Yeast DNA Purification Kit. Bulk pools were sent to the University of Georgia Genomics and Bioinformatics core for KAPA library prep and paired-end 150bp sequencing on an Illumina MiSeq Micro platform for an average coverage of ∼55-fold. DNA from HMY7 was previously sequenced at the Duke Genome Sequencing Core on an Illumina HiSeq 2000 instrument with single-end 50bp reads to an average coverage of 110-fold.

Reads from the bulk pools were aligned to the HMY7 genome (Supplementary Text) using BWA (Li and Durbin 2009), and SNPs were called using Freebayes (Garrison and Marth 2012) with settings for a pooled population. SNPs were filtered for quality and coverage. Bulk pools were compared using the R-package QTLseqr (Mansfeld and Grumet 2018), which implements the smoothed-G statistics of (Magwene et al. 2011).

### YJM311 Subpopulation Membership

In order to assign YJM311 to an *S. cerevisiae* subpopulation, the fixed SNPs from its genome were included in the dataset from the 100-genomes collection and analyzed using the program *structure* V 2.3.4 (Pritchard et al. 2000) following the specifications of Strope et al. (2015). Briefly, the large set of SNPs found across the complete strain panel was filtered and sampled to create four independent sets of ∼1,200 SNPs in low linkage disequilibrium and representing the distribution of minor allele frequencies (generously provided by D. Skelley). Once YJM311 was incorporated, each of the four data sets was run three times using the linkage model (Falush et al. 2003) with a burn-in of 200,000 iterations and 1,000,000 iterations of MCMC, and K=6 groups. The results from the 12 independent runs were compared using *CLUMMP* V 1.1.2 (Jakobsson and Rosenberg 2007) and visualized using *distruct* V 1.1 (Rosenberg 2004).

### Pseudohyphal Growth

For the 100-genomes strains, YPD cultures were grown to saturation (∼24 hours) in a 96-well plate and ∼2 µl per well was transferred to OmniTrays (Nunc 264728) using a 96-pin multi-blot replicator (V&P Scientific no. VP408FP6). For a given assay, a single 96-well plate was pinned to four replicates of three different media types (SLAD, SLAD + PheOH, SLAD + TrpOH). OmniTrays were wrapped with parafilm to prevent drying and incubated at 30C for one week. After incubation, trays were scanned on an Epson Expression 11000 XL scanner, which produced RGB color images with 1200 dpi. For the F5 mapping population, the same procedure was implemented for the 360 segregrants, but only SLAD + PheOH medium was used and with only two replicates per assay. For both the 100-genomes panel and the mapping population, the entire assay was repeated three times.

Follow-up experiments required streaking freezer cultures onto YPD agar, then streaking isolated colonies onto SLAD agar (+ PheOH or TrpOH, when appropriate) and incubating at 30C for 5 days before imaging.

### Image Analysis

The scanned images were processed using a custom Python script (referred to here as “Eclipse”; Supplementary Material) that utilized the skimage package (van der Walt et al. 2014) to read the color qualities of individual pixels. Eclipse discriminated between outer-colony filamentous growth and the inner colony, and reported the ratio of the two, a metric inspired by (Tronnolone et al. 2017).

### Image Processing

It was necessary to identify the color thresholds that designated the colony ring exclusively as white, the filamentous growth as gray, and the background as a separate entity. The image of the entire OmniTray was used to establish the values that best separated the parts of the colony; these values were then used to process the 96 cropped images representing individual colonies. Cracks, smudges, light reflection, and localized contamination interfered with image processing. In these cases, the individual colony images were examined and cropped to exclude trouble spots or dropped from analysis. For YJM311, only segregants that were consistently high-psh and low-psh were of interest for pooling in the bulk segregant analysis. We therefore manually inspected all images and dropped measurements that did not appear to accurately reflect the level of filamentation in the image (assessed qualitatively). This was not done for the panel of environmental isolates because we did not want to introduce bias and because variation in the measurements was of interest for the downstream analysis.

### Statistics

The data from the 100-genomes panel was analyzed in JMP 11.2.0 using an ANOVA framework with three different models. First, no group identity was assigned to the strains. The following model was fitted to the data: Y = µ + Treatment + Strain + Strain* Treatment + Assay + Assay* Treatment + Strain*Assay + Strain*Assay*Treatment. Strain and treatment were considered fixed effects, while assay was considered a random effect. Next, strains were assigned to an ecological niche, which was considered a fixed effect, and the following model was fitted to the data: Y = µ + Niche + Treatment + Strain[Niche] + Assay + Niche*Treatment + Strain* Treatment[Niche] + Assay* Treatment + Assay*Niche + Strain*Assay[Niche] + Strain*Assay*Treatment[Niche]. Brackets denote nested effects. Finally, strains were assigned to a subpopulation and the data were fitted to a model similar to the previous one. The data from the YJM311 segregants were transformed into z-scores for each plate; these values were used to help identify the strains with highest and lowest filamentous index.

### Data Availability

Environmental strains from the 100-genomes collection are available upon request, as well as from the authors of the original study (Strope et al. 2015). Table S1 lists all the isolates along with the filamentous indices extracted from the images. HMY7 and all its segregants are available upon request; table S2 lists all the filamentous indices. Text of the Python program is available as a supplementary file.

All supplementary files are available on figshare. Raw reads for HMY7 and the bulk pools will be deposited in SRA.

## Results

In order to investigate natural variation in psh and its induction by the QS compounds PheOH and TrpOH, the first 96 strains of the 100-genomes panel were plated on nitrogen-limiting medium using a pinning tool. Figure 3 shows the typical structure of a psh colony formed from pinning. Such colonies contain a white “ring” around the inner part where the pinner left cells; this ring separates the grey filamentous growth from the rest of the colony. Our image analysis pipeline located the highest and lowest values of the white ring on both the vertical and horizontal axes, which established a major and minor axis for the ring, and thus mapped out its location as an ellipse. This ellipse was used to “eclipse” all pixels inside of it, demarcating the inner colony. These eclipsed pixels were separated from the non-eclipsed pixels and the ratio of the two was calculated. The ratio, or “filamentous index”, represents a rough quantitative measure of the filamentous growth of the colony; sample values are shown in Figure 3.

**Figure 3:**
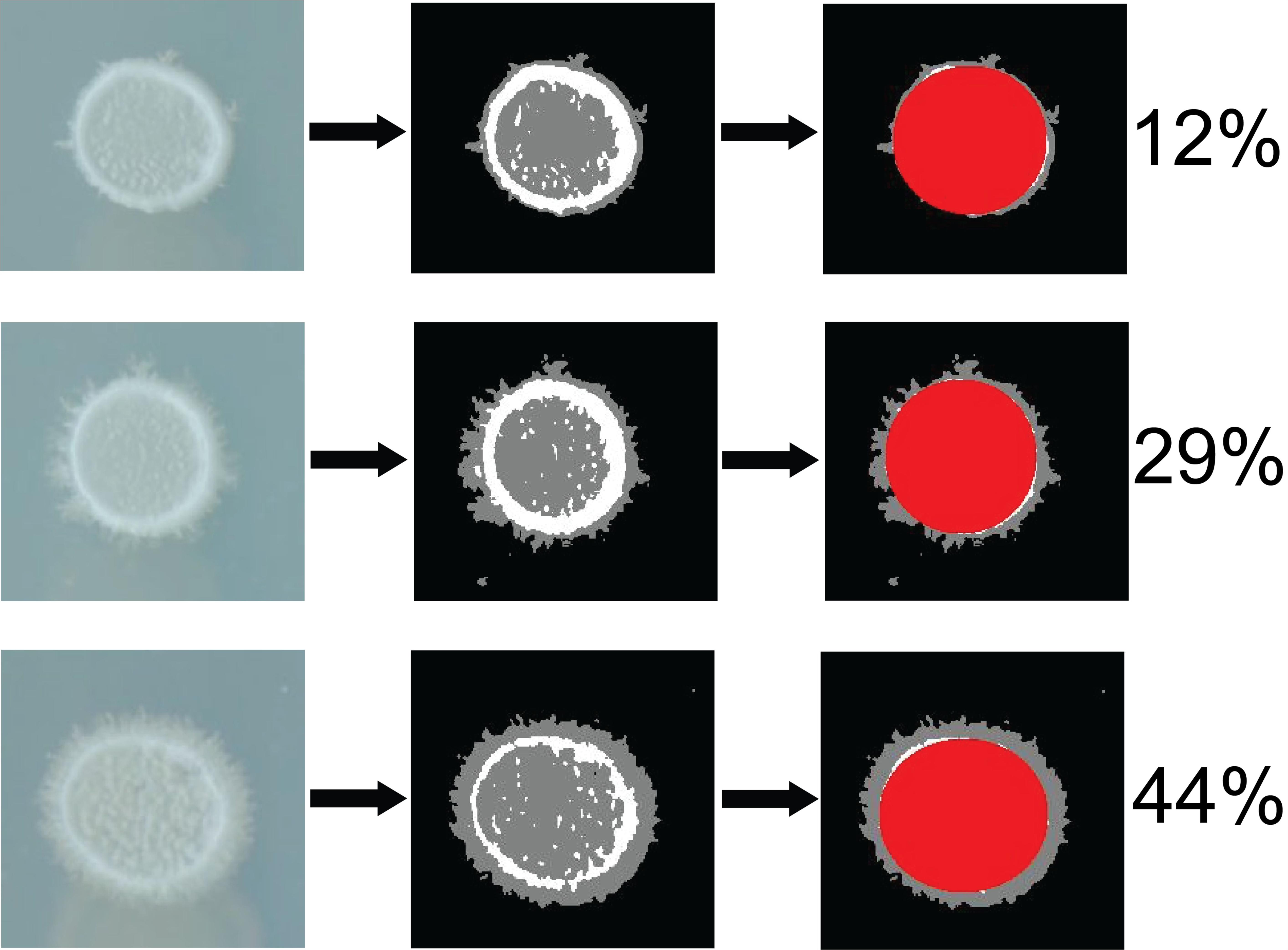
Image processing pipeline. First column: three sample colonies from an Omnitray, which was blotted with 1.58 mm pins; in order, strains YJM984, YJM1336, and YJM1341, derived from 96-112, a clinical strain, M28s2, a European wine strain, and NRRL Y-12637, a South African wine strain, respectively. Second column: original images processed to differentiate white ring, filamentous growth, and background. Third column: inner part of the colony separated and pseudohyphal pixels counted to generate the filamentous index listed on the right.

Across 29 agar trays in three independent assays, 2516 colonies were scored for psh. In the complete data set, the mean filamentous index was 13.01 and the median was 12. These data were analyzed using three linear models. The first did not assign any group identity to the strains and was used to investigate the variation among strains. The second model assigned an ecological niche to the strains and tested for an effect of niche. The third and final model assigned subpopulation membership to the strains and tested for an effect of this phylogenetic history.

### Variation in Pseudohyphal Growth

Of the ∼2500 colonies that were imaged and scored, 895 were grown without the addition of quorum sensing compounds and represent the base level of psh for the strains; the mean filamentous index was 13.29. Overall, there was a wide range of variation in the panel (black data points and distribution in Figure 4A-B) with a maximum average filamentous index of 33.3 for YJM1439 (derived from NCYC110), a ginger beer strain from West Africa, and a minimum average filamentous index of 7.5 for YJM1433 (derived from Yllc17_E5), a wine strain from France.

**Figure 4:**
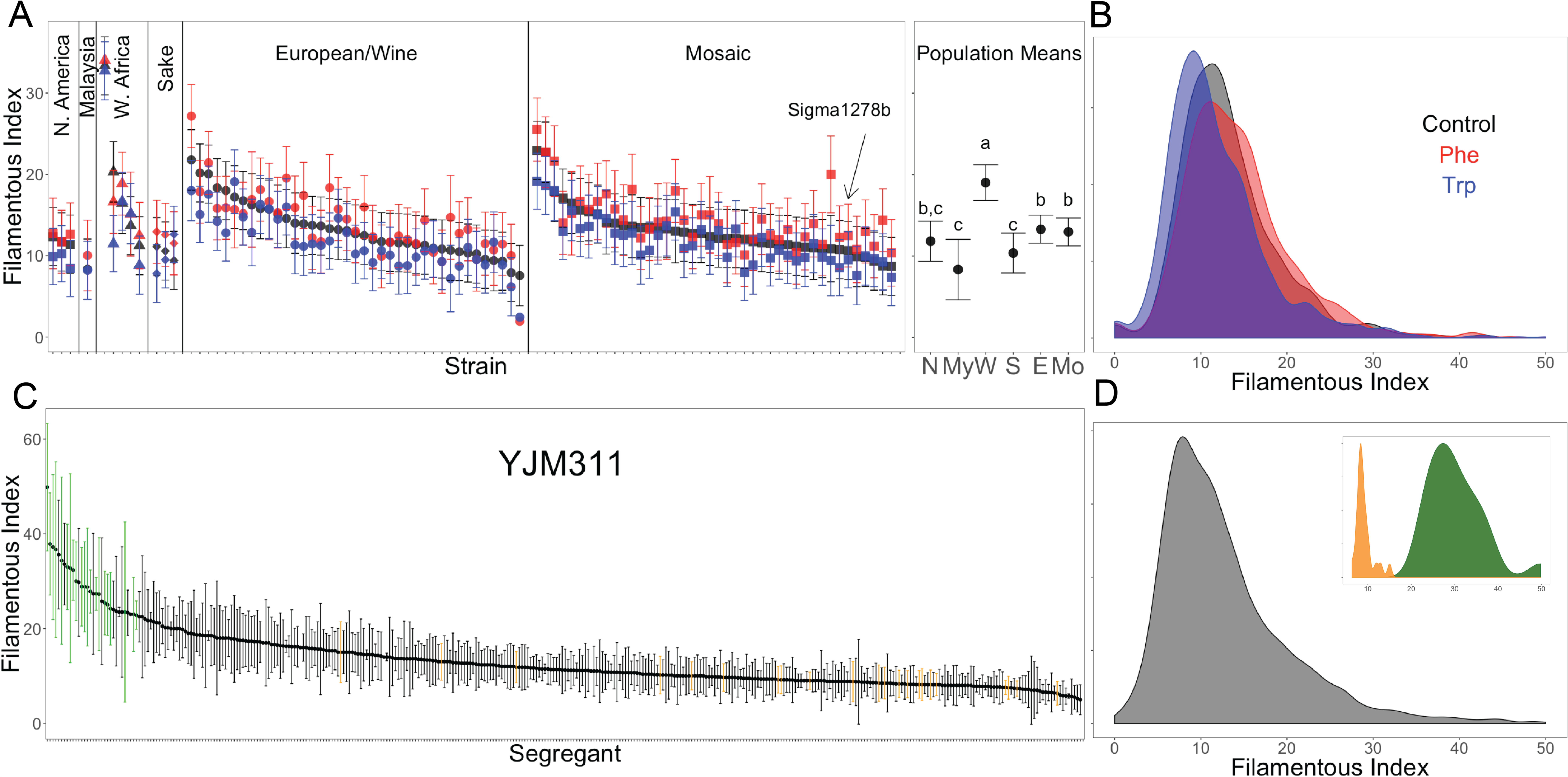
Psh for the 100-genomes panel (A-B), and the YJM311 mapping population (C-D). A and C plot average filamentous index for individual strains or segregants (+/- 2 s.e.m.) and were ordered based on their filamentous index. Panel A also contains the means for the subpopulations; points not connected by the same letter are significantly different. Panels B and D represent population distributions. In A-B, black is control, red is PheOH treatment, and blue is tryptophol treatment. In C-D, green is the high pool and orange is the low pool.

In all three linear models fitted to the data, the strain effect was significant (Table 1). Individual strains with significant parameter estimates (both above and below the mean) are listed in Table 2; strains that were significant in all three models are bolded. While the filamentous index only provides an approximate measure of psh, the behavior of individual strains appears to have been captured well, as the random effects in the model that were associated with replicate assays contributed little of the variation (a total of ∼15% among all the random effects).

**Table 1:** Results of the 100 Genomes pseudohyphal and quorum sensing analyses. Degrees of freedom are estimates due to different numbers of samples in each category and to incomplete samples for some strains (i.e., images dropped from analysis).

| **Model 1: No Group Membership** |  |  |  | 729 |
| --- | --- | --- | --- | --- |
| **Source** | **DF** | **F-ratio** | **p-value** |  |
| Treatment | 2, 4.03 | 2.016 | 0.2473 | 730 |
| Strain | 92, 176.7 | 10.850 | <0.0001 | 731 |
| Treatment*Strain | 184, 343.4 | 0.8994 | 0.7888 | 732 |
| **Random Effects** | **% Total Variance** |  |  | 733 |
| Assay | 0.0 |  |  | 734 |
| Treatment*Assay | 8.9 |  |  | 735 |
| Strain*Assay | 5.4 |  |  | 736 |
| Strain*Assay*Treatment | 0 |  |  | 737 |
| Residual | 85.7 |  |  | 738 |
|  |  |  |  | 739 |
| **Model 2: Niche Membership** |  |  |  | 740 |
| **Source** | **DF** | **F-ratio** | **p-value** |  |
| Niche | 3, 6.33 | 4.35 | 0.0562 | 741 |
| Treatment | 2, 4.41 | 1.946 | 0.2478 | 742 |
| Strain[Niche] | 89, 170.2 | 11.38 | <0.0001 | 743 |
| Treatment*Niche | 6, 358.7 | 0.482 | 0.8217 | 744 |
| Strain*Treatment[Niche] | 178, 343.3 | 0.911 | 0.7574 | 745 |
| **Random Effects** | **% Total Variance** |  |  | 746 |
| Assay | 0.0 |  |  | 747 |
| Treatment*Assay | 8.8 |  |  | 748 |
| Niche*Assay | 1.0 |  |  | 749 |
| Strain*Assay[Niche] | 5.2 |  |  | 750 |
| Strain*Assay*Treatment[Niche] | 0 |  |  | 751 |
| Residual | 85.0 |  |  | 752 |
|  |  |  |  | 753 |
| **Model 3: Population Membership** |  |  |  | 754 |
| **Source** | **DF** | **F-ratio** | **p-value** |  |
| Population | 5, 6.93 | 23.73 | 0.0003 | 755 |
| Treatment | 2, 5.13 | 1.17 | 0.2857 | 756 |
| Strain[Population] | 87, 167.3 | 9.74 | <0.0001 | 757 |
| Treatment*Population | 10, 360.5 | 0.438 | 0.9274 | 758 |
| Strain*Treatment[Pop] | 174, 344.1 | 0.926 | 0.7147 | 759 |
| **Random Effects** | **% Total Variance** |  |  | 760 |
| Assay | 0.0 |  |  | 761 |
| Treatment*Assay | 8.86 |  |  | 762 |
| Population*Assay | 0.03 |  |  | 763 |
| Strain*Assay[Population] | 5.92 |  |  | 764 |
| Strain*Assay*Treatment[Pop] | 0 |  |  | 765 |
| Residual | 85.19 |  |  | 766 |
|  |  |  |  | 767 |

**Table 2:** Parameter estimates for strains that were significant in at least one of the linear models. A p-value of less than 0.05 indicates a significant difference from 0; strains that were significant in all three models are in bold. The parameter estimates are from a linear model and indicate the amount a given strain is above or below the estimate of the intercept. The intercepts for the models are as follows: no group membership-13.16, niche-13.17, population-12.59. In the models with either a niche or population classification, the strains were nested within their group. The estimate for a strain is therefore the combination of the intercept, the strain parameter, and the group parameter.

| Individual Strains |  |  |  |  |  |  |  |  |  |
| --- | --- | --- | --- | --- | --- | --- | --- | --- | --- |
| Well | Genetic Background | Effect Estimate | p-value | Niche | Effect Estimate | p-value | Population | Effect Estimate | p-value |
| G10 | **NCYC110** | 20.21 | <.0001 | Ferment | 19.08 | <.0001 | West African | 11.43 | <.0001 |
| D2 | **NRRL Y-10988** | 9.38 | <.0001 | Clinical | 9.72 | <.0001 | Mosaic | 9.47 | <.0001 |
| F1 | **NRRL Y-12637** | 9.15 | <.0001 | Ferment | 8.02 | <.0001 | European | 9.30 | <.0001 |
| B11 | **R93-1871** | 8.20 | <.0001 | Clinical | 8.52 | <.0001 | Mosaic | 8.30 | <.0001 |
| E11 | **M28s2** | 6.51 | <.0001 | Ferment | 5.38 | <.0001 | European | 6.66 | <.0001 |
| A1 | **YJM128** | 6.02 | <.0001 | Clinical | 6.36 | <.0001 | Mosaic | 6.11 | <.0001 |
| D10 | **NRRL Y-1532** | 4.55 | <.0001 | Plant | 5.17 | <.0001 | European | 4.69 | <.0001 |
| C12 | SK1 | 4.17 | 0.0003 | Lab | 4.32 | <.0001 | West African | -1.14 | 0.2653 |
| A2 | **NCYC 431** | 4.17 | 0.001 | Ferment | 3.04 | 0.0127 | European | 4.31 | 0.0006 |
| H4 | **DBVPG1853** | 3.57 | 0.0018 | Ferment | 2.44 | 0.0255 | European | 3.72 | 0.0011 |
| H5 | **NRRL Y-581** | 3.55 | 0.0016 | Ferment | 2.41 | 0.0249 | European | 3.69 | 0.001 |
| A5 | **CBS 1227** | 2.98 | 0.0083 | Clinical | 3.31 | 0.0025 | European | 3.12 | 0.0053 |
| D11 | NRRL Y-1546 | 2.98 | 0.0086 | Ferment | 1.84 | 0.0895 | West African | -2.34 | 0.0223 |
| A3 | NCYC 914 | 2.89 | 0.0178 | Ferment | 1.78 | 0.1284 | European | 3.032 | 0.0123 |
| F7 | **NRRL Y-11857** | 2.58 | 0.0421 | Plant | 3.22 | 0.0084 | Mosaic | 2.68 | 0.0336 |
| D4 | **MMRL 125** | 2.45 | 0.0398 | Clinical | 2.74 | 0.0177 | Mosaic | 2.54 | 0.0314 |
| F6 | **NRRL Y-5511** | 2.40 | 0.0309 | Plant | 3.02 | 0.0049 | European | 2.55 | 0.0213 |
| D9 | **NRRL Y-35** | 2.40 | 0.0312 | Plant | 3.02 | 0.0049 | European | 2.55 | 0.0215 |
| G1 | NRRL YB-4081 | 2.25 | 0.0532 | Plant | 2.89 | 0.0100 | Mosaic | 2.35 | 0.0425 |
| B1 | B70302(b) | 2.05 | 0.0828 | Clinical | 2.38 | 0.0387 | Mosaic | 2.15 | 0.0673 |
| A4 | NCYC 762 | 1.50 | 0.2336 | Ferment | 0.39 | 0.7461 | West African | -3.81 | 0.0007 |
| G4 | NRRL Y-268 | -1.09 | 0.3258 | Ferment | -2.23 | 0.037 | European | -0.94 | 0.3903 |
| E10 | M1-2 | -1.19 | 0.2852 | Ferment | -2.34 | 0.0302 | European | -1.05 | 0.3437 |
| G5 | NRRL YB-2541 | -1.63 | 0.1426 | Ferment | -2.78 | 0.0098 | European | -1.48 | 0.178 |
| H3 | Y12 | -1.97 | 0.0766 | Ferment | -3.12 | 0.0038 | Sake | 0.29 | 0.7452 |
| A6 | CBS 2910 | -2.24 | 0.0443 | Clinical | -1.91 | 0.0769 | European | -2.09 | 0.0578 |
| B12 | R93-1017 | -2.25 | 0.0447 | Clinical | -1.91 | 0.0792 | Mosaic | -2.16 | 0.053 |
| C4 | 96-101 | -2.26 | 0.0424 | Clinical | -1.92 | 0.0743 | European | -2.11 | 0.0555 |
| F11 | NRRL Y-17447 | -2.28 | 0.0405 | Plant | -1.66 | 0.1193 | Sake | -0.02 | 0.9851 |
| G7 | NRRL YB-2625 | -2.29 | 0.0416 | Plant | -1.68 | 0.1175 | Mosaic | -2.19 | 0.0494 |
| E4 | **Sigma1278b** | -2.29 | 0.0394 | Lab | -2.15 | 0.0169 | Mosaic | -2.20 | 0.0469 |
| A7 | CBS 2807 | -2.30 | 0.0777 | Ferment | -3.41 | 0.007 | European | -2.16 | 0.0951 |
| B8 | **Y55** | -2.31 | 0.0431 | Lab | -2.17 | 0.0172 | West African | -7.62 | <.0001 |
| E2 | YPS134 | -2.34 | 0.036 | Plant | -1.72 | 0.1067 | North American | -0.41 | 0.653 |
| E5 | **RM11** | -2.38 | 0.0324 | Ferment | -3.53 | 0.0011 | European | -2.24 | 0.0427 |
| G12 | UWOPS83-787.3 | -2.38 | 0.0323 | Plant | -1.76 | 0.0978 | Mosaic | -2.29 | 0.0387 |
| D12 | NRRL Y-6673 | -2.40 | 0.0309 | Plant | -1.79 | 0.0932 | European | -2.26 | 0.0409 |
| F4 | **NRRL Y-747** | -2.43 | 0.029 | Ferment | -3.58 | 0.0010 | European | -2.29 | 0.0384 |
| F5 | **NRRL YB-427** | -2.45 | 0.0277 | Ferment | -3.60 | 0.0009 | Mosaic | -2.36 | 0.0332 |
| C5 | **96-109** | -2.50 | 0.0248 | Clinical | -2.16 | 0.0449 | European | -2.36 | 0.033 |
| F10 | NRRL Y-12769 | -2.54 | 0.0238 | Ferment | -3.68 | 0.0008 | Sake | -0.28 | 0.7605 |
| H2 | 273614N | -2.57 | 0.0312 | Clinical | -2.22 | 0.0551 | European | -2.42 | 0.0403 |
| E8 | **UM400** | -2.65 | 0.0174 | Clinical | -2.32 | 0.0319 | Mosaic | -2.56 | 0.021 |
| E7 | **NRRL Y-961** | -2.85 | 0.0106 | Clinical | -2.52 | 0.0198 | Mosaic | -2.76 | 0.0129 |
| G3 | **NRRL YB-4449** | -3.11 | 0.0055 | Plant | -2.50 | 0.0197 | Mosaic | -3.01 | 0.0067 |
| D7 | **UCD-FST 08-200** | -3.13 | 0.0052 | Clinical | -2.79 | 0.0100 | Mosaic | -3.04 | 0.0064 |
| F9 | **NRRL Y-12758** | -3.19 | 0.0044 | Ferment | -4.34 | <.0001 | European | -3.05 | 0.0061 |
| C7 | **96-112** | -3.29 | 0.0034 | Clinical | -2.94 | 0.0066 | European | -3.14 | 0.0047 |
| C3 | **96-100** | -3.37 | 0.0026 | Clinical | -3.03 | 0.0052 | European | -3.23 | 0.0037 |
| F3 | **NRRL Y-234** | -3.40 | 0.0024 | Ferment | -4.55 | <.0001 | European | -3.26 | 0.0034 |
| B9 | **YJM653** | -3.76 | 0.0012 | Clinical | -3.44 | 0.0022 | Mosaic | -3.66 | 0.0015 |
| H1 | **UWOPS05-227.2** | -4.30 | 0.0003 | Plant | -3.68 | 0.0012 | Malaysian | na | na |
| D6 | **UCD-FST 08-199** | -4.38 | 0.0001 | Clinical | -4.04 | 0.0003 | Mosaic | -4.28 | 0.0002 |
| B2 | **B68019c** | -5.13 | <.0001 | Clinical | -4.79 | <.0001 | European | -4.98 | <.0001 |
| G8 | **Yllc17_E5** | -9.18 | <.0001 | Ferment | -10.33 | <.0001 | European | -9.04 | <.0001 |

**Table 2:** Parameter estimates for strains that were significant in at least one of the linear models. A p-value of less than 0.05
|  | Niche | Effect Estimate | p-value | Population | Effect Estimate | p-value |
| --- | --- | --- | --- | --- | --- | --- |
|  | Clinical | -0.328519 | 0.3636 | European | 0.4255585 | 0.1938 |
|  | **Ferment** | 1.1219714 | 0.0151 | **Malaysian** | -3.732617 | 0.0002 |
|  | Lab | -0.155135 | 0.7775 | Mosaic | 0.4744595 | 0.1495 |
|  | Plant | -0.638317 | 0.111 | **North American** | -1.354773 | 0.0249 |
|  |  |  |  | **Sake** | -1.693082 | 0.005 |
|  |  |  |  | **West African** | 5.8804544 | <.0001 |

In the second linear model, strains were divided into four ecological categories based on where they were isolated: fermentation, clinical, plant, and lab environments. The lab category represents strains that have been propagated in the lab environment for many years and may no longer represent the characteristics of the niche from which they were isolated, and includes the model strain Σ1278b. Each of the niche categories contained a wide range of variation in psh (Figure S2). The effect of niche was on the margin of significance in the linear model (*p*=0.056); a *post hoc* Tukey’s Honestly Significant Difference test found fermentation to be higher than the other categories (mean filamentous index of 14.29 compared to 13.01, 12.83, and 12.52 for lab, clinical, and plant, respectively). However, if the strain with the most abundant pseudohyphae, YJM1439, is removed from the analysis, the niche effect is no longer significant (*p*=0.185; fermentation mean = 13.50), suggesting the effect is tenuous.

In the third and final linear model, strains were assigned membership to a subpopulation (based on the *structure* analysis) (Figures 4A,B). Most of the strains fell in the European/wine and mosaic categories, with the Malaysian subpopulation represented by a single strain; therefore the results for this analysis should be interpreted with caution. The effect of population was significant in the model (*p*=0.0003). A *post hoc* Tukey’s Honestly Significant Difference test found the West African subpopulation to have a higher filamentous index than the other categories (Figure 4B, last panel). The West African subpopulation contained YJM1439, the strain with the highest filamentous index. When this strain was removed, the West African subpopulation remained significantly higher than the others (mean=14.77 compared to 18.47 with YJM1439). Thus, for at least one subpopulation, membership may be an important predictor for psh.

### Variation in Response to Quorum Sensing Compounds

Of the colonies that were imaged and scored, 670 were grown in medium supplemented with PheOH and 951 were grown in medium supplemented with TrpOH; these produced a mean filamentous index of 14.20 and 11.89 respectively. Surprisingly, there was no overall effect of the addition of QS compounds (*p*=0.2473 in the model with no group identity, *p*=0.2478 in the niche model, and *p*=0.2857 in the population model). Two strains significantly increased psh in response to PheOH in all three models and a different two strains increased in response to TrpOH. However, two strains also significantly *decreased* psh in the presence of one or both of the compounds (Table 3). Even for the strains that appeared to respond significantly, the effect sizes were small (on the order of 2-3 in the filamentous index). Mostly, PheOH and TrpOH appeared to have little to no effect on psh.

**Table 3:** Parameter estimates for response to the QS treatments. A p-value of less than 0.05 indicates a significant difference from 0; strains that were significant in all three models are in bold. Strains whose phenotypic response was investigated via streaking have a yes or no to indicate whether the predicted response was detectable; “-” indicates the strain was not streaked.

| Individual Strain x Treatment Effects |  |  |  |  |  |  |  |  |  |  |
| --- | --- | --- | --- | --- | --- | --- | --- | --- | --- | --- |
| Well | Verified | Treatment | Effect Estimate | p-value | Niche | Effect Estimate | p-value | Population | Effect Estimate | p-value |
| **D11** | **yes** | **Control** | 4.08 | 0.0008 | Ferment | 4.23 | 0.0005 | West African | 3.64 | 0.001 |
| **G8** | – | **Control** | 3.48 | 0.0049 | Ferment | 3.65 | 0.0029 | European | 3.31 | 0.0071 |
| D10 | no | Control | 2.37 | 0.0429 | Plant | 2.21 | 0.0554 | European | 2.20 | 0.0588 |
| F6 | – | Control | 2.34 | 0.0454 | Plant | 2.18 | 0.0588 | European | 2.17 | 0.0622 |
| E9 | – | Control | 2.28 | 0.051 | Clinical | 2.24 | 0.0539 | Mosaic | 2.48 | 0.0335 |
| G12 | no | Control | -2.12 | 0.0694 | Plant | -2.29 | 0.0472 | Mosaic | -1.93 | 0.0963 |
| H5 | **yes** | Control | -2.24 | 0.0559 | Ferment | -2.09 | 0.0709 | European | -2.41 | 0.038 |
| **A9** | **yes** | **Phe** | 5.34 | 0.0009 | Clinical | 5.15 | 0.0012 | Mosaic | 5.20 | 0.0011 |
| **F1** | no | **Phe** | 3.62 | 0.0046 | Ferment | 3.66 | 0.0039 | European | 3.76 | 0.0031 |
| D1 | no | Phe | 2.63 | 0.0474 | Clinical | 2.52 | 0.056 | European | 2.76 | 0.0359 |
| G12 | no | Phe | 2.35 | 0.0591 | Plant | 2.63 | 0.0321 | Mosaic | 2.20 | 0.0754 |
| **G8** | – | **Phe** | -3.27 | 0.0106 | Ferment | -3.26 | 0.0099 | European | -3.14 | 0.0135 |
| **A2** | **yes** | **Trp** | 3.08 | 0.0236 | Ferment | 2.89 | 0.0306 | European | 3.13 | 0.0206 |
| **H3** | no | **Trp** | 2.75 | 0.016 | Ferment | 2.57 | 0.023 | Sake | 2.36 | 0.0124 |
| G11 | no | Trp | 2.29 | 0.0452 | Plant | 2.17 | 0.0536 | Mosaic | 2.24 | 0.0481 |
| **F1** | no | **Trp** | -3.00 | 0.0154 | Ferment | -3.18 | 0.0092 | European | -2.95 | 0.0161 |
| **D11** | **yes** | **Trp** | -3.34 | 0.0039 | Ferment | -3.53 | 0.002 | West African | -3.17 | 0.0026 |

**Table 3:** Parameter estimates for response to the QS treatments.
| Treatment Effects |  |  |  |  |  |  |  |  |
| --- | --- | --- | --- | --- | --- | --- | --- | --- |
| Treatment | Effect Estimate | p-value | Treatment | Effect Estimate | p-value | Treatment | Effect Estimate | p-value |
| Control | 0.095 | 0.9034 | Control | 0.023 | 0.9773 | Control | 0.040 | 0.9608 |
| Phe | 1.223 | 0.1694 | Phe | 1.268 | 0.1588 | Phe | 1.194 | 0.1854 |
| Trp | -1.318 | 0.1455 | Trp | -1.290 | 0.1526 | Trp | -1.234 | 0.171 |

### Comparison to Streaked Colonies

In order to verify the results from the high throughput assay, a selection of 10 strains that appeared to respond significantly to the QS molecules were streaked on SLAD, SLAD + PheOH, and SLAD + TrpOH agar plates (Figure S3). We qualitatively assessed whether there appeared to be more filamentation in the different treatments (Table 3) in a manner similar to the study that originally reported the effects of the QS molecules on Σ1278b (Chen et al. 2006). We found that 4 of the 10 strains appeared to respond in the direction predicted, as best as could be detected from visual inspection, but all responses were subtle.

The colonies arising from the streaks on SLAD agar were also compared to the images of the pinned colonies on the SLAD OmniTrays in order to verify that the high throughput method was correctly assessing the overall status of psh ability. While more psh was induced via streaking than pinning, there was clear agreement between the methods: strains with strong psh in one method exhibited strong psh in the other, while non-psh strains did not produce filamentation in either method (Figure S3). However, it is also clear that filamentous index is a rough measurement, as strains that had similar psh induction did not have precisely the same index values. This is likely because all colonies stemming from one OmniTray were analyzed with the same color thresholds. This approach was taken in order to avoid bias, but future work analyzing each colony with its own optimized thresholds could potentially make the filamentous index more accurate. As it is currently being implemented, it appears to be appropriate for assessing general relative psh ability in a large panel.

### Natural Genetic Variation in Pseudohyphal Growth

In order to investigate the amount of natural segregating genetic variation for psh, an F5 mapping population of YJM311 was phenotyped, and high and low segregants were pooled for sequencing and analysis. Across 24 agar trays, 360 segregants produced 1823 colonies that were scored. The range of phenotypic variation within the mapping population was comparable to that of the 100-genomes collection of environmental strains (Figure 4B,D): the segregants had an overall mean filamentous index of 13.03 with a median of 11, and the maximum and minimum average filamentous index values were 49.8 and 5, respectively. The pools of segregants used in the sequencing analysis had distinct phenotypic distributions with a high pool mean of 30.32, (standard deviation= 6.45), and a low pool mean of 9.13 (standard deviation = 1.96) (Figure 3D, insert).

### Bulk Segregant Analysis

The allele frequencies of the bulk pools were compared using a smoothed-G statistic in order to find chromosomal regions that contain variation associated with psh (Magwene et al. 2011, Mansfeld et al. 2018). Different window sizes for the smoothing function generated a variable number of significant mapping peaks, with windows of 60KB, 40KB, and 20KB producing 4, 26, and 29 significant peaks, respectively, at a false discovery rate of 0.01 (Figure 5). While a smaller window size is likely to be appropriate for an F5 mapping population, we highlighted candidate genes in the four peaks stemming from the 60KB window, as well as one more on chromosome 14 which just missed the cut-off, as these peaks likely represent major effect loci (Table 4). In all five peaks, there were numerous genes that have been shown to either increase or decrease pseudohyphal or invasive growth in the Σ1278b background.

**Table 4:** Candidate genes from the bulk segregant analysis listed with chromosome, general function, and whether or not there is published data linking the gene to filamentous growth in Σ1278b. For each gene, variation separating the high and low bulks in YJM311 is listed by the number of SNPs in the coding region (synonymous or non-synonymous), and the number of SNPs occurring 300 base pairs upstream and/or downstream of the coding region (possible regulatory variation). 1-(Kang et al. 2005), 2-(Shively et al. 2013), 3-(Guo et al. 2000), 4-(Mosch and Fink 1997), 5-(Jin et al. 2008), 6- (Song et al. 2014), 7-(Laxman and Tu 2011), 8-(Zhu et al. 2000), 9-(Scherz et al. 2014), 10-(Lorenz et al. 1998), *-includes SNPs in intron.

| Chr | Gene | Function | Association in $\Sigma 1278b$ | Syn. | Non-Syn. | Reg. ($\pm 300bp$) |
| --- | --- | --- | --- | --- | --- | --- |
| 3 | ABP1 | transcription factor | Yes<sup>1,2</sup> | 2 | 1 | 2 |
|  | FIG2 | cell adhesin | Yes<sup>3</sup> | 11 | 18 | 7 |
|  | CDC39 | transcriptional regulator | Yes<sup>4</sup> | 24 | 10 | 3 |
| 5 | PEA2 | polarisome subunit | Yes<sup>1,5,6</sup> | 2 | 0 | 1 |
|  | SPI1 | cell wall protein | Yes<sup>5</sup> | 2 | 0 | 12 |
| 13 | AIM33 | protein of unknown function | No | 3 | 0 | 6 |
|  | ALO1 | catalytic enzyme | Yes<sup>2</sup> | 5 | 0 | 3 |
|  | YML083C | protein of unknown function | No | 2 | 1 | 1 |
|  | YML082W | putative protein | No | 2 | 3 | 0 |
|  | WAR1 | transcription factor | Yes<sup>2</sup> | 0 | 5 | 0 |
| 14 | APJ1 | chaperone | Yes<sup>2,5</sup> | 0 | 5 | 3 |
|  | MKS1 | transcriptional regulator | Yes<sup>2,5,7</sup> | 0 | 0 | 1 |
|  | MSK1 | mito. tRNA synthetase | Yes<sup>5</sup> | 1 | 7 | 4 |
|  | FKH2 | transcription factor | Yes<sup>8</sup> | 1 | 8 | 5 |
|  | SUN4 | cell wall protein | Yes<sup>2,5</sup> | 0 | 5 | 11 |
|  | GCD10 | tRNA methyltransferase | Yes<sup>5</sup> | 0 | 2 | 16 |
| 16 | RPS23B | ribosomal protein | No, but see<sup>1</sup> | 1 | 0 | 8* |
|  | TOM5 | outer membrane translocase | Yes<sup>9</sup> | 0 | 0 | 6 |
|  | MEP3 | ammonium permease | Yes<sup>10</sup> | 3 | 0 | 3 |
|  | KAR3 | microtubule motor | Yes<sup>2</sup> | 2 | 3 | 1 |
|  | ASN1 | Asparagine synthetase | No | 16 | 3 | 6 |
|  | YPR148C | protein of unknown function | Yes<sup>2</sup> | 8 | 3 | 2 |

**Figure 5:**
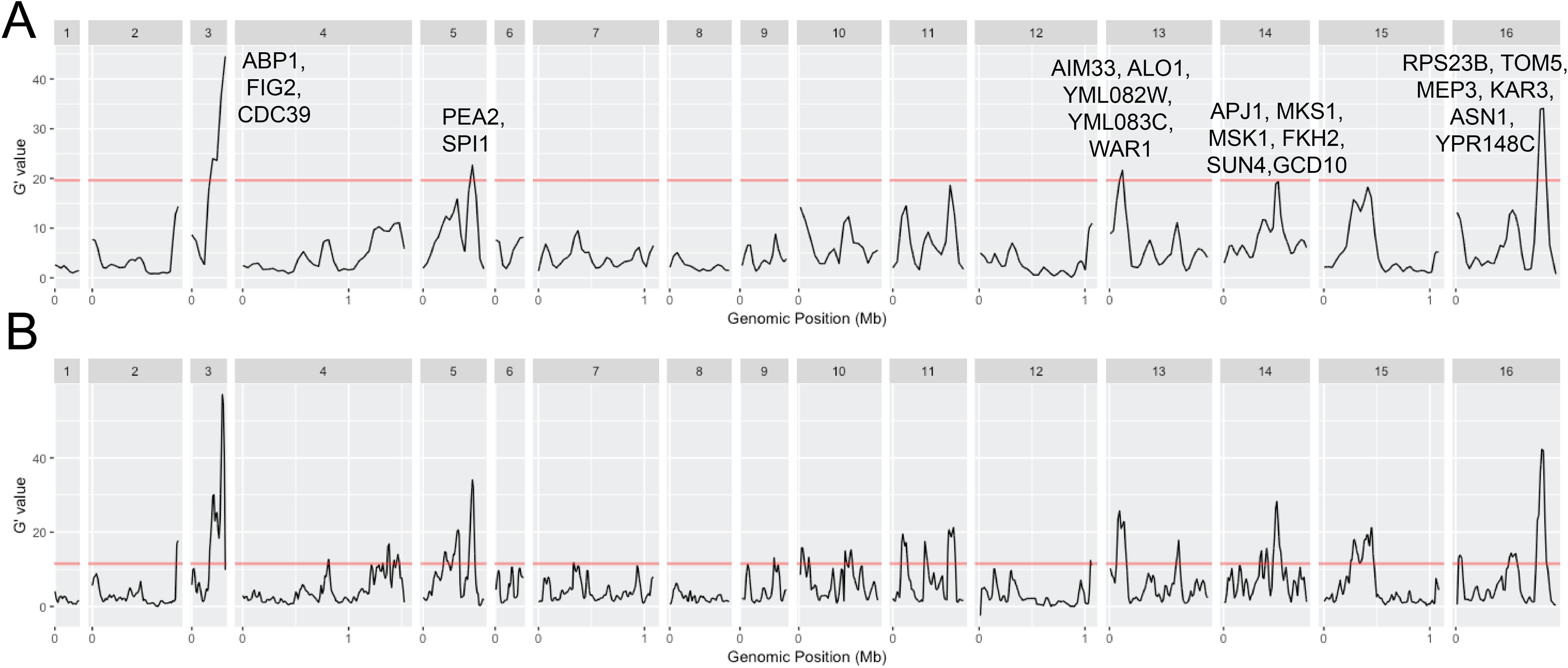
Genome-wide comparison of the allele frequencies in the high and low psh pools of YJM311 F5 segregants. The G-prime statistic was calculated with a sliding window size of 60,000 bp (A) and 20,000 bp (B). Red line represents the cut-off for significance at a false discovery rate of 0.01. Candidate loci are listed above major, significant peaks.

## Discussion

Microbes can engage in a myriad of social phenotypes that provide fitness benefits to individuals and genetic lineages (West et al. 2006). The model yeast, *S. cerevisiae*, exhibits multiple social phenotypes in the lab, including filamentous growth and quorum sensing. The filamentous growth phenotype appears to be conserved among other *Saccharomyces spp* (Kayikci and Magwene 2018) and among medically relevant yeasts, including *Candida albicans* (Cutler 1991), other *Candida spp* (Silva et al. 2011), *Asperigillus fumigatus* (Mowat et al. 2009) and *Trichosporon asahii* (Di Bonaventura et al. 2006), with filamentation ranging from pseudohyphae to true hyphae. Thus, filamentous growth is likely an important fungal response to environmental cues. This phenotype can be induced via QS in both *S. cerevisiae* (Chen et al. 2006) and *C. albicans* (Hornby et al. 2001, Chen et al. 2004), though the QS molecules are not shared.

The goal of this study was to assess the extent of variation in psh and response to external QS molecules in a range of isolates of *S. cerevisiae* in order to understand how the traits vary in natural populations. As such, we were interested in a strong response that is robust to slight environmental fluctuations. Our experimental protocol differed from those used in other studies of psh: we did not use Noble agar (highly purified) or wash cells before plating, and we attempted to quantify the amount of pseudohyphae rather than agar invasion. We also pinned from culture, transferring thousands of cells, rather than streaking to generate colonies from a single cell; our observation is that psh is more extensive when a colony is generated from a single cell (Figure S3). The relative consistency among methods, assays, and replicate plates suggests our results captured the phenotype well, and represents an estimate of the general filamentous response in these environmental strains.

### Phenotypic Variation in Pseudohyphal Growth

There was a surprising amount of phenotypic variation in the 100-genomes collection, with some strains exhibiting long, pronounced pseudohyphae and some strains having little to no pseudohyphal growth. The image analysis counted a small amount of the white ring of the colony; therefore, values below 10% represent no, or very little, psh. It is possible that if these low strains were assayed in a different manner (i.e., streaked on highly processed medium) more psh would be observed; however, our goal was to assay a general, robust psh response and these strains lacked one.

We hypothesized that clinical strains would exhibit a stronger phenotype due to the association of filamentous growth with biofilm formation and virulence in other yeasts (Fanning and Mitchell 2012). However, when the strains were divided into their ecological niche of origin, there did not appear to be a particular niche that had more psh than others. Filamentous growth is likely a more general response and the perceived association with virulence may simply be due to a bias in the organisms in which research is conducted. Another possibility, which is not mutually exclusive, is that *S. cerevisiae* is not adapted to specific ecological niches with regard to this phenotype. Rather, as was previously proposed by (Goddard and Greig 2015), it is a “nomad” dispersed among many habitats due to its association with humans. Our analysis that divided strains by their population of origin suggested that certain subpopulations are associated with increased psh, specifically, the West African subpopulation. This supports the idea that there is a signal based on phylogeny rather than ecology for this trait.

### Phenotypic Variation in Response to QS Molecules

Based on research in Σ1278b, we hypothesized that some of the low-psh strains would be induced when PheOH and TrpOH were added exogenously. We also hypothesized that fermentation strains would be most sensitive to the QS compounds, as the molecules could disperse further in more viscous environments where QS could be beneficial in synchronizing populations. Furthermore, a recent study investigated the effect of compounds produced during aromatic amino acid metabolism on different wine yeast (González et al. 2018). Non-*Saccharomyces* yeast growth was negatively affected by the presence of TrpOH and PheOH, suggesting that these compounds could be particularly important in inter-species interactions in fermentation environments.

Surprisingly, most strains in the 100-genomes collection did not respond to the addition of QS molecules to the medium. It is still possible that many of these strains use PheOH and TrpOH for communication, but that the response is too subtle to be detected in our assay (see below). However, if cells do indeed carefully regulate both the production of and the response to QS compounds, it is improbable that exogenous application would have so little detectable effect across a wide panel. At the very least, one would expect a slight change in the same direction in most strains, which is not what we observed. Instead, it is more likely that certain strains respond strongly to PheOH and TrpOH, but most simply do not.

### Comparison to Σ1278b

The majority of research on filamentous growth and QS in *S. cerevisiae* has been done on strains derived from Σ1278b, which has proved an invaluable model for understanding the genetic basis of the trait and for generating a robust map of the genetic pathways controlling it (Cullen et al. 2012). Homologs of some of the genes implicated in the Σ1278b background have been shown to be important for filamentous growth in other yeast species (Lo et al. 1997, Cain et al. 2012). And in the present study, genes uncovered in Σ1278b potentially harbor causative allelic variation in the clinical isolate YJM311.

Chen and Fink (2006) demonstrated changes in the amount of pseudohyphae produced when Σ1278b was exposed to dilute treatments of PheOH, TrpOH, and both in combination. This strain was included in our panel (well E4), and while it appeared to somewhat respond to one of the autoinducing chemicals (PheOH), our results were not as dramatic as theirs. This is likely because our phenotypic assay was not as sensitive: our analysis measures “fuzziness” around a large colony, so the difference between treatments has to be striking to be detected. When we streaked, rather than pinned, Σ1278b, our results were similar to the previously published results (Figure S4), but the amount of change induced is small compared to the range of variation found among environmental isolates. In our assay, we did find individual strains that significantly responded to both chemicals, as was expected. However, in the majority of strains, the results were not as anticipated, and in some cases, were actually the opposite of Σ1278b. It is possible that other strains in our panel could also have a subtle response to the QS compounds, but it is clear that in most strains, the molecules do not induce a dramatic phenotypic change. The difference in inducibility between Σ1278b and the majority of strains in the panel indicates a disparity in behavior between this popular model strain and environmental strains of *Saccharomyces cerevisiae*. Thus, when it comes to QS, the results from the model laboratory strain may not translate smoothly to the broader population of *Saccharomyces* yeasts and how they behave in the environment.

### Genetic Variation

The present study aimed to determine whether and how much natural allelic variation existed in psh by using in a heterozygous clinical isolate from the mosaic subpopulation as a proxy. The phenotypic variation in the mapping population recapitulated the variation in the environmental panel, and five major and many minor peaks were associated with the trait, suggesting an abundance of segregating variation for psh in the environment. Complex phenotypes can be strongly influenced by SNPs at non-synonymous, synonymous, and regulatory locations (She and Jarosz 2018); all these types of genetic variation were identified in the major mapping peaks of YJM311. We mostly highlighted candidate genes in the peaks that have been implicated in psh in the Σ1278b background, but it is not clear whether or not they contain the causative alleles. These loci influence numerous cellular processes such as cell wall biosynthesis, mitochondrial function (Kang and Jiang 2005), cell polarity (Song et al. 2014), progression through the cell cycle (Zhu et al. 2000), and ammonium uptake (Lorenz and Heitman 1998). While investigating the functional effect of various alleles was beyond the scope of this study, we anticipate future studies could harness the power of this approach.

It is worth noting that a recent study comparing psh in *S. cerevisiae* and *S. bayanus* found that the cyclic AMP-Protein Kinase A pathway plays an important regulatory role in both. However, the manner in which the genetic network regulates the phenotype has diverged: increasing levels of cAMP has the opposite effect on the induction of the phenotype in the two species (Kayikci et al. 2018). This suggests selection to maintain filamentous growth over a long time scale, but also the ability of the complex genetic network underlying the trait to adapt and change. Future work identifying the genetic basis of some of the phenotypic variation observed in this study could shed light on the components of the genetic network that currently harbor segregating allelic variation, and upon which selection could ultimately act.

## Acknowledgements

We thank Paul Magwene for strains, Daniel Skelley for genomic data files, Rachel Rambadt for computational assistance, and the University of Georgia Genomics and Bioinformatics core for sequencing the samples. The research was funded by National Institutes of Health grant R15GM122032 to H.A.M. and a William & Mary Biology DeFontes Fellowship to B.A.L.

## References Cited

Biswas, S., P. Van Dijck and A. Datta (2007). Environmental Sensing and Signal Transduction Pathways Regulating Morphopathogenic Determinants of *Candida albicans*. Microbiology and Molecular Biology Reviews 71(2): 348.

Braus, G. H., O. Grundmann, S. Bruckner and H. U. Mosch (2003). Amino acid starvation and Gcn4p regulate adhesive growth and FLO11 gene expression in Saccharomyces cerevisiae. Mol Biol Cell 14(10): 4272–4284.

Cain, C. W., M. B. Lohse, O. R. Homann, A. Sil and A. D. Johnson (2012). A Conserved Transcriptional Regulator Governs Fungal Morphology in Widely Diverged Species. Genetics 190(2): 511–521.

Chen, H. and G. R. Fink (2006). Feedback control of morphogenesis in fungi by aromatic alcohols. Genes Dev 20(9): 1150–1161.

Chen, H., M. Fujita, Q. Feng, J. Clardy and G. R. Fink (2004). Tyrosol is a quorum-sensing molecule in *Candida albicans*. Proceedings of the National Academy of Sciences of the United States of America 101(14): 5048.

Chen, R. E. and J. Thorner (2010). Systematic Epistasis Analysis of the Contributions of Protein Kinase A- and Mitogen-Activated Protein Kinase-Dependent Signaling to Nutrient Limitation-Evoked Responses in the Yeast Saccharomyces cerevisiae. Genetics 185(3): 855–870.

Cullen, P. J. and G. F. Sprague (2012). The Regulation of Filamentous Growth in Yeast. Genetics 190(1): 23–49.

Cullen, P. J. and G. F. Sprague, Jr. (2000). Glucose depletion causes haploid invasive growth in yeast. Proc Natl Acad Sci U S A 97(25): 13619–13624.

Cutler, J. E. (1991). Putative Virulence Factors of Candida Albicans. Annual Review of Microbiology 45(1): 187–218.

Di Bonaventura, G., A. Pompilio, C. Picciani, M. Iezzi, D. D’Antonio and R. Piccolomini (2006). Biofilm formation by the emerging fungal pathogen Trichosporon asahii: development, architecture, and antifungal resistance. Antimicrob Agents Chemother 50(10): 3269–3276.

Falush, D., M. Stephens and J. K. Pritchard (2003). Inference of population structure: Extensions to linked loci and correlated allele frequencies. Genetics 164: 1567–1587.

Fanning, S. and A. P. Mitchell (2012). Fungal Biofilms. PLoS Pathog 8(4): e1002585.

Garrison, E. and G. Marth (2012). Haplotype-based variant detection from short-read sequencing. arXiv 1207.3907.

Gasch, A. P., B. A. Payseur and J. E. Pool (2016). The power of natural variation for model organism biology. Trends in Genetics 32: 146–154.

Gimeno, C. J., P. O. Ljungdahl, C. A. Styles and G. R. Fink (1992). Unipolar cell divisions in the yeast S. cerevisiae lead to filamentous growth: regulation by starvation and RAS. Cell 68(6): 1077–1090.

Goddard, M. R. and D. Greig (2015). Saccharomyces cerevisiae: a nomadic yeast with no niche? FEMS Yeast Research 15(3): fov009–fov009.

González, B., J. Vázquez, P. J. Cullen, A. Mas, G. Beltran and M.-J. Torija (2018). Aromatic Amino Acid-Derived Compounds Induce Morphological Changes and Modulate the Cell Growth of Wine Yeast Species. Frontiers in Microbiology 9(670).

Granek, J. A., Ö. Kayikçi and P. M. Magwene (2011). Pleiotropic signaling pathways orchestrate yeast development. Current Opinion in Microbiology 14(6): 676–681.

Granek, J. A., D. Murray, Ö. Kayikçi and P. M. Magwene (2013). The Genetic Architecture of Biofilm Formation in a Clinical Isolate of *Saccharomyces cerevisiae*. Genetics 193(2): 587–600.

Guo, B., C. A. Styles, Q. Feng and G. R. Fink (2000). A Saccharomyces gene family involved in invasive growth, cell-cell adhesion, and mating. Proc Natl Acad Sci U S A 97(22): 12158–12163.

Hornby, J. M., E. C. Jensen, A. D. Lisec, J. J. Tasto, B. Jahnke, R. Shoemaker, et al. (2001). Quorum Sensing in the Dimorphic Fungus *Candida albicans* Is Mediated by Farnesol. Applied and Environmental Microbiology 67(7): 2982.

Jakobsson, M. and N. Rosenberg (2007). *CLUMMP*: a cluster matching and permuation program fro dealing with label switching and mulitmodality in analysis of population structure. Bioinformatics 23: 1801–1806.

Jin, R., C. J. Dobry, P. J. McCown and A. Kumar (2008). Large-scale analysis of yeast filamentous growth by systematic gene disruption and overexpression. Mol Biol Cell 19(1): 284–296.

Kang, C. M. and Y. W. Jiang (2005). Genome-wide survey of non-essential genes required for slowed DNA synthesis-induced filamentous growth in yeast. Yeast 22(2): 79–90.

Kayikci, Ö. and P. M. Magwene (2018). Divergent Roles for cAMP–PKA Signaling in the Regulation of Filamentous Growth in *Saccharomyces cerevisiae* and *Saccharomyces bayanus*. G3: Genes|Genomes|Genetics 8(11): 3529.

Lambrechts, M. G., F. F. Bauer, J. Marmur and I. S. Pretorius (1996). Muc1, a mucin-like protein that is regulated by Mss10, is critical for pseudohyphal differentiation in yeast. Proc Natl Acad Sci U S A 93(16): 8419–8424.

Laxman, S. and B. P. Tu (2011). Multiple TORC1-associated proteins regulate nitrogen starvation-dependent cellular differentiation in Saccharomyces cerevisiae. PLoS One 6(10): e26081.

Leberer, E., D. Harcus, D. Dignard, L. Johnson, S. Ushinsky, D. Y. Thomas, et al. (2001). Ras links cellular morphogenesis to virulence by regulation of the MAP kinase and cAMP signalling pathways in the pathogenic fungus Candida albicans. Molecular Microbiology 42(3): 673–687.

Li, H. and R. Durbin (2009). Fast and accurate long-read alignment with Burrows-Wheeler Transform. Bioinformatics 25: 1754–1760.

Liti, G., D. M. Carter, A. M. Moses, J. Warringer, L. Parts, S. A. James, et al. (2009). Population genomics of domestic and wild yeasts. Nature 458: 337–341.

Lo, H. J., J. R. Kohler, B. DiDomenico, D. Loebenberg, A. Cacciapuoti and G. R. Fink (1997). Nonfilamentous C. albicans mutants are avirulent. Cell 90(5): 939–949.

Lo, W.-S. and A. M. Dranginis (1998). The Cell Surface Flocculin Flo11 Is Required for Pseudohyphae Formation and Invasion by Saccharomyces cerevisiae. Molecular Biology of the Cell 9(1): 161–171.

Lorenz, M. C. and J. Heitman (1998). Regulators of pseudohyphal differentiation in Saccharomyces cerevisiae identified through multicopy suppressor analysis in ammonium permease mutant strains. Genetics 150(4): 1443–1457.

Magwene, P. M., Ö. Kayikçi, J. A. Granek, J. M. Reininga, Z. Scholl and D. Murray (2011). Outcrossing, mitotic recombinatino, and life-history trade-offs shape genome evolution in *Saccharomyces cerevisiae*. Proceedings of the National Academy of Sciences of the USA 108(5): 1987–1992.

Magwene, P. M., J. H. Willis and J. K. Kelly (2011). The Statistics of Bulk Segregant Analysis Using Next Generation Sequencing. PLoS Comput Biol 7(11): e1002255.

Mallick, E. M. and R. J. Bennett (2013). Sensing of the Microbial Neighborhood by <italic>Candida albicans</italic>. PLoS Pathog 9(10): e1003661.

Mansfeld, B. N. and R. Grumet (2018). QTLseqr: An R Package for Bulk Segregant Analysis with Next-Generation Sequencing. The Plant Genome 11(2).

Matsui, T., R. Linder, J. Phan, F. Seidl and I. M. Ehrenreich (2015). Regulatory Rewiring in a Cross Causes Extensive Genetic Heterogeneity. Genetics 201(2): 769.

Mosch, H. U. and G. R. Fink (1997). Dissection of filamentous growth by transposon mutagenesis in Saccharomyces cerevisiae. Genetics 145(3): 671–684.

Mowat, E., C. Williams, B. Jones, S. McChlery and G. Ramage (2009). The characteristics of Aspergillus fumigatus mycetoma development: is this a biofilm? Med Mycol 47 Suppl 1: S120–126.

Pan, X. and J. Heitman (1999). Cyclic AMP-dependent protein kinase regulates pseudohyphal differentiation in Saccharomyces cerevisiae. Mol Cell Biol 19(7): 4874–4887.

Pritchard, J., M. Stephens and P. Donnelly (2000). Inference of population structure using multilocus genotypes. Genetics 155: 945–959.

Replansky, T., V. Koufopanou, D. Greig and G. Bell (2008). *Saccharomyces sensu stricto* as a model system for evolution and ecology. Trends in Ecology and Evolution 23(9): 494–501.

Rocha, C. R., K. Schroppel, D. Harcus, A. Marcil, D. Dignard, B. N. Taylor, et al. (2001). Signaling through adenylyl cyclase is essential for hyphal growth and virulence in the pathogenic fungus Candida albicans. Mol Biol Cell 12(11): 3631–3643.

Rosenberg, N. (2004). *Distruct*: a program for the graphical display of population structure. Molecular Ecology Notes 4: 137–138.

Rupp, S., E. Summers, H. J. Lo, H. Madhani and G. Fink (1999). MAP kinase and cAMP filamentation signaling pathways converge on the unusually large promoter of the yeast FLO11 gene. Embo Journal 18(5): 1257–1269.

Ryan, O., R. S. Shapiro, C. F. Kurat, D. Mayhew, A. Baryshnikova, B. Chin, et al. (2012). Global gene deletion analysis exploring yeast filamentous growth. Science 337(6100): 1353–1356.

Scherz, K., Andersen, R. Bojsen, L. Gro, Rejkjaer, Sorensen, et al. (2014). Genetic basis for Saccharomyces cerevisiae biofilm in liquid medium. G3 (Bethesda) 4(9): 1671–1680.

She, R. and D. F. Jarosz (2018). Mapping Causal Variants with Single-Nucleotide Resolution Reveals Biochemical Drivers of Phenotypic Change. Cell 172(3): 478-490.e415.

Shively, C. A., M. J. Eckwahl, C. J. Dobry, D. Mellacheruvu, A. Nesvizhskii and A. Kumar (2013). Genetic Networks Inducing Invasive Growth in Saccharomyces cerevisiae Identified Through Systematic Genome-Wide Overexpression. Genetics 193(4): 1297–1310.

Silva, S., M. Negri, M. Henriques, R. Oliveira, D. W. Williams and J. Azeredo (2011). Adherence and biofilm formation of non-Candida albicans Candida species. Trends in Microbiology 19(5): 241–247.

Song, Q., C. Johnson, T. E. Wilson and A. Kumar (2014). Pooled Segregant Sequencing Reveals Genetic Determinants of Yeast Pseudohyphal Growth. PLoS Genet 10(8): e1004570.

Strope, P. K., D. A. Skelly, S. G. Kozmin, G. Mahadevan, E. A. Stone, P. M. Magwene, et al. (2015). The 100-genomes strains, an S. cerevisiae resource that illuminates its natural phenotypic and genotypic variation and emergence as an opportunistic pathogen. Genome Research 25(5): 762–774.

Tronnolone, H., J. M. Gardner, J. F. Sundstrom, V. Jiranek, S. G. Oliver and B. J. Binder (2017). Quantifying the dominant growth mechanisms of dimorphic yeast using a lattice-based model. Journal of The Royal Society Interface 14(134).

van der Walt, S., J. L. Schönberger, J. Nunez-Iglesias, F. Boulogne, J. D. Warner, N. Yager, et al. (2014). scikit-image: image processing in Python. PeerJ 2: e453.

Warringer, J., E. Zörgö, F. A. Cubillos, A. Zia, A. Gjuvsland, J. T. Simpson, et al. (2011). Trait Variation in Yeast Is Defined by Population History. PLOS Genetics 7(6): e1002111.

West, S. A., A. S. Griffin, A. Gardner and S. P. Diggle (2006). Social evolution theory for microorganisms. Nat Rev Micro 4(8): 597–607.

Wuster, A. and M. M. Babu (2010). Transcriptional control of the quorum sensing response in yeast. Mol Biosyst 6(1): 134–141.

Zaman, S., S. I. Lippman, X. Zhao and J. R. Broach (2008). How Saccharomyces responds to nutrients. Annu Rev Genet 42: 27–81.

Zhu, G., P. T. Spellman, T. Volpe, P. O. Brown, D. Botstein, T. N. Davis, et al. (2000). Two yeast forkhead genes regulate the cell cycle and pseudohyphal growth. Nature 406(6791): 90–94.

